# Ramping activity in midbrain dopamine neurons signifies the use of a cognitive map

**DOI:** 10.1101/2020.05.21.108886

**Authors:** Akash Guru, Changwoo Seo, Ryan J. Post, Durga S. Kullakanda, Julia A. Schaffer, Melissa R. Warden

**Author notes:** NYU Grossman School of Medicine, New York University, 550 1st Ave, New York, NY 10016. Laboratory of Behavioral and Genomic Neuroscience, National Institute on Alcohol Abuse and Alcoholism, Rockville, MD 20852, USA.

## Abstract

Journeys to novel and familiar destinations employ different navigational strategies. The first drive to a new restaurant relies on map-based planning, but after repeated trips the drive is automatic and guided by local environmental cues^1,2^. Ventral striatal dopamine rises during navigation toward goals and reflects the spatial proximity and value of goals^3^, but the impact of experience, the neural mechanisms, and the functional significance of dopamine ramps are unknown^4,5^. Here, we used fiber photometry^6–8^ to record the evolution of activity in midbrain dopamine neurons as mice learned a variety of reward-seeking tasks, starting recordings before training had commenced and continuing daily for weeks. When mice navigated through space toward a goal, robust ramping activity in dopamine neurons appeared immediately – after the first rewarded trial on the first training day in completely naïve animals. In this task spatial cues were available to guide behavior, and although ramps were strong at first, they gradually faded away as training progressed. If instead mice learned to run a fixed distance on a stationary wheel for reward, a task that required an internal model of progress toward the goal, strong dopamine ramps persisted indefinitely. In a passive task in which a visible cue and reward moved together toward the mouse, ramps appeared and then faded over several days, but in an otherwise identical task with a stationary cue and reward ramps never appeared. Our findings provide strong evidence that ramping activity in midbrain dopamine neurons reflects the use of a cognitive map^9,10^ – an internal model of the distance already covered and the remaining distance until the goal is reached. We hypothesize that dopamine ramps may be used to reinforce locations on the way to newly-discovered rewards in order to build a graded ventral striatal value landscape for guiding routine spatial behavior.

---

The decision to continue pursuing a goal or abandon the quest depends on how much progress has been made, how much remains to be done, and the value of the goal. For example, a climber will be more deterred by rain at a mountain’s base than near the summit, and will be more reluctant to abandon a prized peak than a training hill. Information about progress toward goals and their value is essential for adaptively balancing time and energy between activities, and commitment to goals and the vigour of goal-directed actions are both regulated by goal progress and value^11–16^.

Ventral striatal dopamine (DA) progressively rises as rodents navigate toward spatially distant rewards^3^, a surprising recent finding that was not anticipated by temporal difference learning models of DA function^4^ but which has broadened our understanding of the role of ventral striatal DA in sustaining and invigorating goal-directed behavior^17^. DA ramps reflect the value and proximity of goals, scaling by the value of the reward and stretching or compressing in different environments to span the distance between start and goal locations^3^. Ventral striatal DA also rises during self-initiated actions to obtain rewards^18–23^, and ramping activity has been observed in ventral tegmental area (VTA) DA neurons via single-unit electrophysiology and calcium imaging^24–30^. These observations have inspired efforts to understand the functional significance of DA ramps, which have been hypothesized to signal reward expectation, reward prediction error, discounted reward, state value, or uncertainty^3–5,23,30–36^. Here, we sought to probe the neural mechanisms underlying DA ramps by examining how ramping activity in VTA DA neurons evolves with experience and how this evolution is affected by the presence or absence of informative sensory cues.

We began by asking whether activity in VTA DA neurons increases as animals navigate through space toward goals, as ventral striatal DA release can occur independently of VTA DA neural activity^31,37–40^. The VTA of DAT-Cre mice^41^ was injected with a Cre-dependent genetically encoded calcium indicator, AAV-DJ-EF1α-DIO-GCaMP6m^42^, and an optical fiber was implanted over the VTA (Fig. 1a) in order to record population activity in DA neurons via fiber photometry^6–8^. Mice learned a simple self-paced reward-seeking task, in which they ran back and forth on a long linear track to obtain a ‘big’ reward (10 µL of water) at one end and a ‘small’ reward (5 µL) at the other (Fig. 1b). Activity in DA neurons increased as mice ran toward rewards (Fig. 1c-f, Extended Data Table 1; GFP control data, Extended Data Fig. 1), and continuously rising activity was evident on individual trials (Fig. 1c, d). DA neuron activity did not reflect instantaneous run speed, which peaked near the middle of the track (Extended Data Fig. 2a, b). Ramp slope depended on reward magnitude (Fig. 1e, f) and followed the locations of the big and small rewards as they switched positions (Extended Data Fig. 2c). Surprisingly, DA ramps on the linear track emerged immediately – on the first day of training in completely naïve animals – during the first run to a goal location following a single experience with reward (Fig. 1g-k). We observed this phenomenon for runs to both the big reward and runs to the small reward (Extended Data Fig. 3). This rapid appearance of DA ramps is unlike the timescale of cue-triggered DA reward prediction error (RPE) signals, which emerge slowly after hundreds of conditioning trials^43,44^, but is reminiscent of the timescale of fast hippocampal remapping^45–48^.

**Fig. 1.**
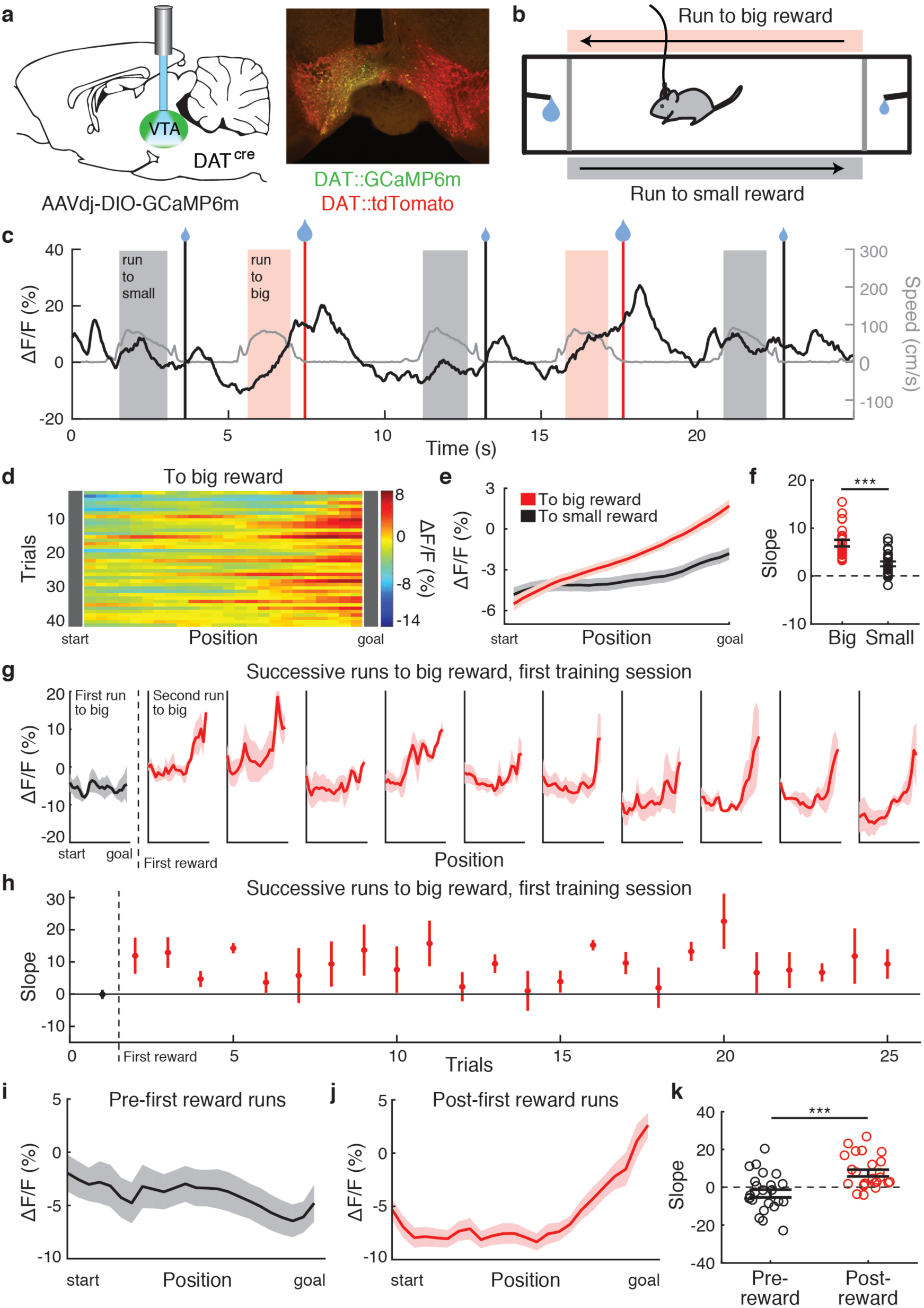
Ramping activity in VTA DA neurons appears immediately after the first-ever rewarded trial in naïve mice. **a**, Recording configuration, left. GCaMP6m expression in VTA DA neurons in a DAT-Cre/Ai14 mouse, right. **b**, Track schematic. **c**, Example track photometry data from a DAT::GCaMP6m mouse. GCaMP ΔF/F in black, speed in grey. **d**, Example ΔF/F heatmap from a DAT::GCaMP6m mouse. **e**, Mean ΔF/F with respect to position for big and small rewards (n=23; 6 animals, 3-4 sessions). **f**, Mean ramp slope during runs to big and small rewards. \*\*\**P* < 0.001, Wilcoxon signed-rank test **g**, Mean ΔF/F for successive runs to the big reward during the first training session. Black – before the first-ever reward exposure; red – after the first-ever reward exposure (n=3). **h**, Ramp slope for successive runs to the big reward during the first track session. **i**, Mean ΔF/F with respect to position for the 8 track runs to the big reward location immediately preceding the first-ever reward delivery (n=24; 3 animals, 8 trials). **j**, Mean ΔF/F with respect to position for the 8 track runs to the big reward location immediately following the first-ever reward delivery (n=24; 3 animals, 8 trials). **k**, Mean ramp slopes for the 8 trials before and after the first-ever reward (n=24; 3 animals, 8 trials). \*\*\**P* < 0.001, Wilcoxon rank-sum test. Error bars indicate s.e.m.

On the track, and in other instrumental tasks where DA ramps have been observed, animals move to obtain rewards. However, ramping activity in DA neurons has been observed during Pavlovian conditioning under some conditions^24^, suggesting that DA ramps may emerge even in the absence of goal-directed movement. To test this idea, we designed a ‘passive approach’ task in which the mouse did not go to the reward – instead, the reward came to the mouse (Fig. 2a). Mice initiated trials by breaking an infrared beam positioned next to a transparent wall with a small opening. A cue light and a reward spout were co-mounted on an actuator positioned 10 cm from the opening, and upon trial initiation the cue light illuminated and the cue/reward began moving toward the mouse. When the cue/reward reached the mouse, the reward could be consumed through the opening. We found that VTA DA neural activity ramped up as the distance between the mouse and the cue/reward decreased, even though no action was required during cue/reward movement (Fig. 2b-e). If the cue/reward was instead stationary and permanently located on the chamber wall, DA ramps were absent (Fig. 2d, e). Thus, observing the progress of an approaching reward was sufficient to produce ramping activity in DA neurons – goal-directed movement was not required.

**Fig. 2.**
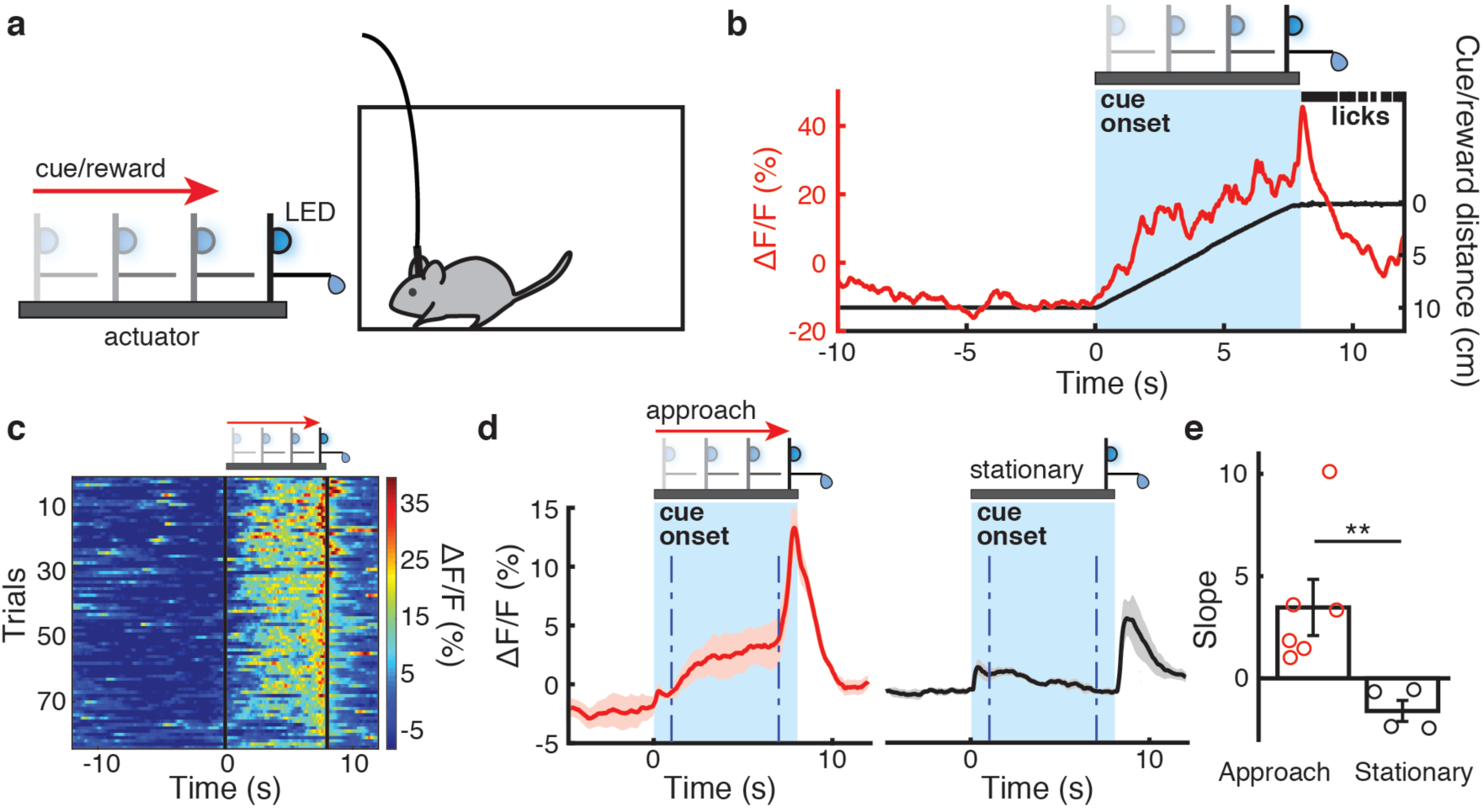
Approaching but not stationary conditioned cues produce VTA DA ramps. **a**, Passive conditioning with approaching cue/reward. **b**, Approaching cue/reward photometry example trial from a DAT::GCaMP6m mouse. GCaMP ΔF/F in red, cue/reward distance in black, licks in black. **c**, Example ΔF/F heat map, approaching cue/reward. **d**, Mean ΔF/F aligned to cue onset for approaching cue/reward (n=6, left) and for stationary cue/reward (n=4, right). Dashed lines indicate slope analysis epoch. **e**, Mean ramp slopes for approaching and stationary cues/rewards. \*\**P <* 0.01, Wilcoxon rank-sum test. Error bars indicate s.e.m.

In the tasks that we have examined thus far, mice could use systematically changing sensory cues to assess their progress toward the goal. We asked whether an internal representation of goal progress was sufficient to produce DA ramps in the absence of these cues. We addressed this question by training mice to run on a wheel for a fixed distance. Mice self-initiated a trial by starting to run, and if they ran without stopping for at least 5 turns of the wheel and stopped before 9 turns, a cue light was illuminated and a water reward was delivered via a spout adjacent to the wheel (‘contingent’ task, Fig. 3a, Extended Data Fig. 4). We also ran a yoked control group in which running was permitted but not required, and in which rewards were delivered on a schedule determined by the contingent group (‘non-contingent’ task, Fig. 3b, Extended Data Fig. 4). During performance of the ‘contingent’ task, DA neural activity robustly increased as mice ran, despite the absence of informative sensory cues (Fig. 3c, e, g; Extended Data Fig. 5a), and ramps were evident on single trials (Fig. 3c). This finding shares some features with internally-generated hippocampal episode/time fields, which have been observed during wheel running^49–51^. During the ‘non-contingent’ task, performed by a separate group of mice, DA neural activity did not increase during running bouts but instead rose steeply upon cue onset/reward delivery (Fig. 3d, f, g; Extended Data Fig. 5b). Thus, an internal model of progress toward the goal is sufficient for the expression of DA ramps – informative sensory cues are not required.

**Fig. 3.**
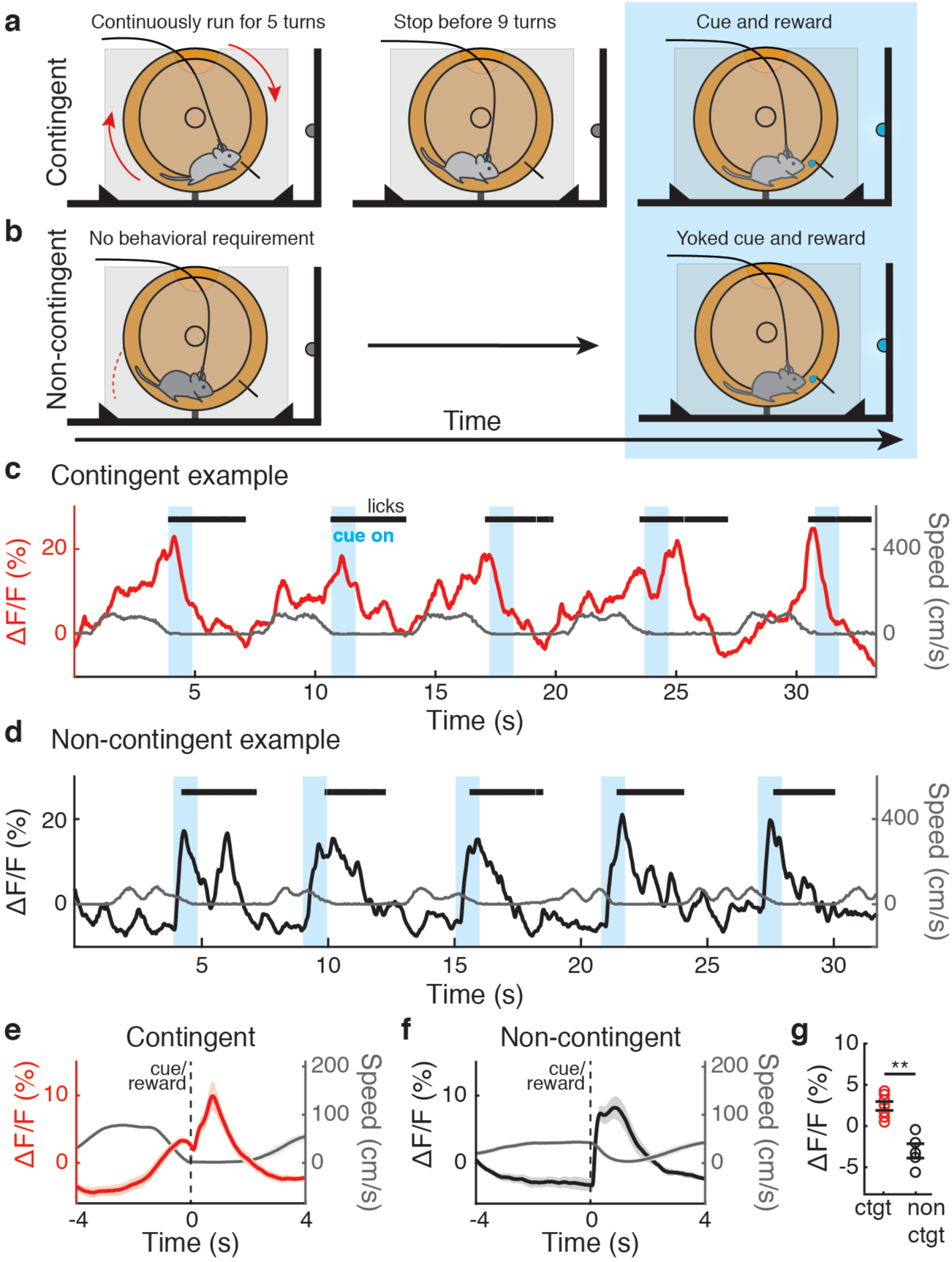
An internal model of progress toward a goal is sufficient to produce VTA DA ramps. **a**, Contingent wheel task schematic. **b**, Non-contingent wheel task schematic. **c**, Example wheel photometry data from a contingent DAT::GCaMP6m mouse. GCaMP ΔF/F in red, speed in grey, licks in black. **d**, Example wheel photometry data from a non-contingent DAT::GCaMP6m mouse. GCaMP ΔF/F in black, speed in grey, licks in black. **e**, Mean ΔF/F (red) and mean speed (grey) aligned to stop-triggered cue/reward onset, contingent group (n=7). **f**, Mean ΔF/F (black) and mean speed (grey) aligned to cue/reward onset, non-contingent group (n=5). **g**, Mean pre-cue ΔF/F, contingent and non-contingent groups. \*\**P <* 0.01, Wilcoxon rank-sum test. Error bars indicate s.e.m.

Finally, we examined the evolution of DA ramps over extended training in all tasks. Ramps on the linear track were steepest on the first and second training days, and gradually faded as daily training continued (Fig. 4a, b). Switching the spatial positions of big and small rewards reinvigorated ramps (Fig. 4a, b), as observed previously^21^, but did not affect run time or path length (Extended Data Fig. 6). DA ramps during passive conditioning with an approaching cue/reward took several days to develop, and gradually transitioned to cue-driven activity as training progressed (Fig. 4c, d). Ramps were never observed during passive conditioning with a stationary cue/reward (Fig. 4c, d). Unlike DA ramps during track runs and passive conditioning, DA ramps on the wheel never faded, remaining robust even over extended training (Fig. 4e, f). The wheel is unique among the tasks that we tested because all systematically changing sensory cues have been eliminated. Thus, it is not possible to develop an association between a unique sensory state and reward, and mice must instead rely on an internal model of progress toward the goal. Comparing the evolution of DA ramps in different tasks reveals that DA ramps persist indefinitely when an internal representation of progress toward the goal is required, but fade when changing sensory cues are available to guide behavior.

**Fig. 4.**
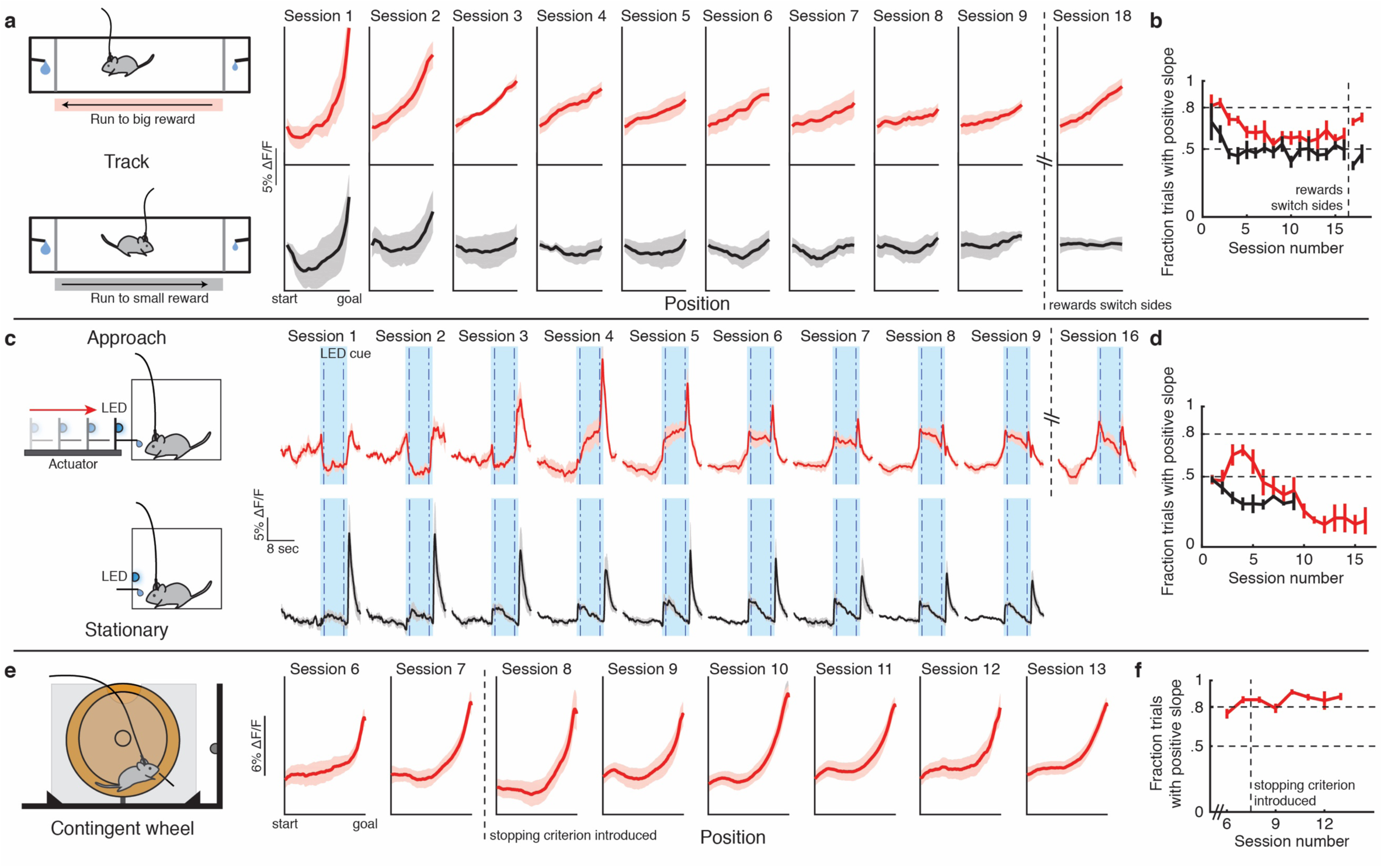
VTA DA ramps persist when an internal model of goal proximity is required but fade when environmental cues are available. **a**, Track schematic, left. Mean ΔF/F with respect to position for track runs to big (top) and small (bottom) rewards by session in DAT::GCaMP6m mice, right (n=3 session 1, n=4 session 2, n=5 sessions 3-18). Rewards switch sides after session 16. **b**, Mean fraction of trials with positive ramp slope for track runs to big (red) and small (black) rewards by session. **c**, Approaching and stationary conditioned cue/reward schematic, left. Mean ΔF/F for approaching (top, n=6 session 1-10, n=3 session 11-16) and stationary (bottom, n=4 session 1-6,8,9, n=3 session 7) cues/rewards by session, right. **d**, Mean fraction of trials with positive ramp slope for approaching (red) and stationary (black) cues/rewards by session. **e**, Wheel schematic, left. Mean ΔF/F with respect to position for wheel runs by session, right (n=6). **f**, Mean fraction of trials with positive ramp slope for wheel runs by session. Error bars indicate s.e.m.

Here, we have examined how ramping activity in VTA DA neurons evolves with experience, and how this evolution is affected by environmental cues. We have shown that DA ramps appear immediately when naïve mice engage in spatial navigation – on the first trial following the initial discovery of reward. We have also shown that DA ramps gradually fade when changing sensory cues are available to guide behavior, but never fade when mice must use an internal model of goal progress. Finally, we have shown that DA ramps don’t require physical movement, but can instead be produced by cues/rewards moving toward passive mice.

Our findings strongly support the hypothesis that ramping activity in dopamine neurons reflects the use of an internal model – a cognitive map of how much progress has been made toward a goal and how much remains to be done – and as such imply that DA ramps reflect goal-directed or model-based behavioral control rather than habitual or model-free^1,52,53^. Our findings cast doubt on the hypothesis that DA ramps reflect the reward prediction error term in temporal difference (TD) learning algorithms. First, there simply isn’t any time for the required repeated sampling and updating. The phasic DA RPE signal^54^ requires hundreds of trials to emerge^43,44^, but our findings reveal that DA ramps appear after a single rewarded trial. Second, TD learning models predict that DA ramps should flatten when systematically changing sensory cues are not available and be maintained when they are available^36^, but here we show the opposite – strong DA ramps persist indefinitely in the absence of informative sensory cues but gradually fade when they are available. These divergent findings and predictions suggest that distinct neural mechanisms likely underlie DA ramps and DA RPE signals.

We speculate that DA ramps may reflect the transmission of goal proximity information from cortical or cortical-like brain regions, such as the hippocampus/entorhinal cortex or orbitofrontal cortex, to VTA DA neurons. The hippocampus is thought to represent a cognitive map of spatial and non-spatial relationships^9,10,55–57^, sends a massive projection to the ventral striatum^58–60^, and regulates VTA DA neural activity via the ventral striatum^61,62^. In addition to dorsal hippocampal place cell sequences that may reflect prospection and retrospection^63–68^, the distance to goals and objects is explicitly encoded in the average firing rates of ventral hippocampus and entorhinal neurons^69–73^. Intriguingly, goal distance encoding in human hippocampus and entorhinal cortex is suppressed when subjects are instructed to use local sensory cues and avoid thinking about goals or directions to goals^74^. Frontal cortical regions including the orbitofrontal cortex (OFC) have also been implicated in encoding goal proximity, and the OFC has been hypothesized to represent a cognitive map of task space^53^. The OFC sends a direct projection to VTA DA neurons^75^, and OFC lesions degrade reward expectancy signals in VTA DA neurons^26^, suggesting a potential role for OFC circuits in the construction of DA ramps. Consistent with this idea, OFC value signals are amplified when animals direct their gaze closer to conditioned cues^76^.

How might DA ramps be used to shape learning? In laboratory environments, stimulation of VTA DA neurons robustly reinforces both place occupancy and actions at specific places^77–79^, but it is unknown whether DA ramps are reinforcing, and if so what they reinforce. One possibility is spatial location, as suggested by the importance of DA in conditioned place preference and the strength of hippocampal inputs to the ventral striatum^58–60^. By reinforcing locations encountered on the way to newly discovered rewards, dopamine ramps could build a spatial value landscape in the ventral striatum, an idea that may generalize to more abstract relational domains^9^. Such a representation could be used as a heuristic to guide routine navigation and other habitual spatial behavior^80–82^, freeing cortical and cortical-like circuits for planning and other effortful, deliberative cognitive operations^83^

## Methods

### Mice

All procedures conformed to guidelines established by the National Institutes of Health and have been approved by the Cornell University Institutional Animal Care and Use Committee. DAT-Cre mice (The Jackson Laboratory, Bar Harbor, ME) were crossed with Ai14 Cre reporter mice (The Jackson Laboratory, Bar Harbor, ME) to generate DAT-Cre/Ai14 mice. DAT-Cre/Ai14 or DAT-Cre mice (postnatal 2-10 months) were used for DA-specific viral vector expression. All Cre driver lines were fully backcrossed to C57BL/6J mice. Both male and female mice were used. All mice were maintained on a 12-hour reverse light-dark cycle with *ad libitum* access to food and water, except during behavioral experiments.

### Viral vectors

We used AAV-DJ-EF1α-DIO-GCaMP6m (Stanford Vector Core, Stanford, CA) or AAV5-CAG-Flex-GFP (UNC Vector Core, Chapel Hill, NC) for photometry experiments.

### Surgery

Mice were anaesthetized using isoflurane (5%) and prepared for stereotactic surgery. Fur was shaved and mice were placed in a stereotaxic frame (Kopf Instruments, Tujunga, CA). A heating pad was placed under the mice to prevent hypothermia. Isoflurane (1-2%) was delivered via a nose cone throughout the surgery. Ophthalmic ointment was used to protect the eyes. Buprenorphine (0.05 mg/kg, subcutaneous) was given before the start of surgery. A mixture of 0.5% lidocaine and 0.25% bupivacaine (100 µL) was injected intradermally along the incision line. The scalp was disinfected with betadine scrub and 70% ethanol. An incision was made using a scalpel along the scalp midline. The exposed skull was thoroughly cleaned, and a craniotomy was made above the VTA.

Virus was targeted to the VTA (−3.1 AP, 0.3-0.4 ML, 4.6 & 4.3 DV), and slowly pressure-injected (100 nl/min) using a 10 µL Hamilton syringe (nanofil; WPI, Sarasota, FL), a 33-gauge bevelled needle, and a micro-syringe pump controller (Micro 4; WPI, Sarasota, FL). After each injection, the needle was left in place for 10 minutes and then slowly withdrawn. A total of 800 nl (500 nl at −4.6 DV and 300 nl at −4.3 DV) of vector was injected. Animals were implanted with a 400 µm diameter, 0.48 NA optical fiber (Doric Lenses, Québec, Canada) above the VTA (−3.1 AP, 0.3-0.4 ML, 4.3-4.4 DV). A layer of metabond (Parkell, Inc., Edgewood, NY) and dental acrylic (Lang Dental Manufacturing, Wheeling, IL) was applied to firmly hold the fiber in place, and the surrounding skin was sutured. Post-operative buprenorphine (0.05 mg/kg), carprofen (5 mg/kg), and lactated ringers (500 µl) were administered subcutaneously. Behavioral testing began after a minimum 3-week viral expression period.

### Perfusion and Histological Verification

Animals were deeply anaesthetized with Fatal-Plus at a dose of 90 mg/kg and transcardially perfused with 20 ml of PBS (phosphate-buffered saline), followed by 20 ml of 4% paraformaldehyde solution. Brains were quickly extracted and stored in 4% paraformaldehyde solution at 4°C for 24 hrs. Brains were then transferred to 30% sucrose in PBS solution and allowed to equilibrate for 2 days. Brains were then mounted on a freezing microtome using OCT and sectioned coronally (45 µm). Sections were washed in PBS and mounted on slides with PVA-DABCO. Images were acquired using a Zeiss LSM 800 confocal scanning laser microscope with a 5X and 20X air objective.

### Fiber Photometry

Fiber photometry was performed as previously described^8^. 473 nm and 405 nm data were collected for all GCaMP and GFP photometry experiments using a custom-built fiber photometry system. 473 nm and 405 nm diode lasers (Omicron Luxx, Rodgau-Dudenhofen, Germany) were modulated at 700 Hz and 500 Hz, respectively, using a dual optical chopper (New Focus Model 3502, Newport, Irvine, CA). The laser beams were combined using a mirror (KM100-E02, Thorlabs, Newton, NJ) and a dichroic filter (LM01-427, Semrock, Rochester, NY), and the power was adjusted using a neutral density filter to 15-80 µW. The laser beams were then aligned to a fluorescence filter cube (DFMB, Thorlabs, Newton, NJ). Within the cube, the excitation beams were reflected with a dichroic filter (FF495, Semrock, Rochester, NY), collimated (F240FC-A with AD11F, Thorlabs, Newton, NJ), and coupled to an optical patch cord (400 µm, Doric Lenses, Quebec, Canada). The optical patch cord was connected to the animal using a zirconia sleeve (ADAF1, Thorlabs, Newton, NJ). Emitted fluorescence received through the same optical patch cord was passed through a dichroic filter (FF495, Semrock, Rochester, NY) and a bandpass filter (FF03-525/50, Semrock, Rochester, NY), and was collected by a femtowatt photoreceiver (New Focus 2151, Newport, Irvine, CA). Voltage from the photoreceiver was channelled through two lock-in amplifiers (SR810 DSP, Stanford Research System, Sunnyvale, CA), digitized using a DAQ (U6 pro, LabJack, Lakewood, CL) at a sampling rate of 250 Hz, and recorded by LabJack software.

### Behavioral Testing

Mice were water restricted prior to all behavioral testing. Body mass was measured daily to maintain at least 80% of baseline body weight.

#### Linear track

A 122 × 12.5 cm custom-made plexiglass track contained reward ports on both ends. Licking was monitored using dual contact lickometers (ENV-250B, Med Associates, Fairfax, VT) when mice completed the circuit between the reward spout and a copper plate placed beneath the reward port. Water rewards were delivered upon lick initiation using variable-rate infusion syringe pumps (PHM-107, Med Associates, Fairfax, VT) on both ends. Mice were required to alternate between the reward ports to trigger reward delivery. 10 µl of water was delivered at one port and 5 µl of water was delivered at the other port. The location of the big reward (10 µl) and the small reward (5 µl) was held constant for a given session but was sometimes switched between sessions, as indicated. Behavioral testing was conducted under dark/red light and each session lasted 20-30 min per day. Video was recorded using an infrared (IR) USB 3.0 camera (Chameleon3, Pointgrey, Richmond, BC, Canada), which provided TTL pulses for every recorded frame. Lick timestamps, reward delivery timestamps, and video TTL pulses were sent to the photometry DAQ for synchronized recording. Video data were processed offline using Ethovision-XT (Noldus Information Technology, VA) to collect position and velocity data. Reward port access was blocked at the beginning of each session for 5 minutes while mice were allowed to freely explore the track and the photometry signal was recorded. For the linear track experiments in Figure 4, a linear track (132 × 12.5 cm) was modified with IR sensors and controllers (ENV-253SD, ENV-253 Med Associates, Fairfax, VT). The IR sensors were placed 22 cm from the reward port on both sides and rewards were delivered when the mouse exited the reward port and crossed the closest IR sensor. The locations of big (8 µl) and small (4 µl) rewards were kept constant for 16 sessions and switched for session 17 and 18. Photometry data was recorded during each session.

#### Running wheel

The behavioral apparatus consisted of a running wheel (Comfort Exercise Wheel, Kaytee Products Inc., Chilton, WI) mounted on a breadboard (Thorlabs, Newton, NJ). The breadboard was kept flat or at an angle of 20 degrees to facilitate running. The wheel was connected to an optical encoder (E4T 200 cycles per revolution, US Digital, Vancouver, WA) using a custom-made shaft to record wheel movement. A blue LED visual cue was mounted in front of the wheel. Clear plexiglass covered the front of the wheel. A metal lick spout was attached to the plexiglass and was connected to a custom capacitive lickometer. Water reward was delivered using a syringe pump (NE-500, New Era Pump Systems, Farmingdale, NY). All hardware operation and behavioral data acquisition (1 KHz sampling rate) were controlled by a data acquisition card (USB-6212, National Instruments, Austin, TX) with custom LabVIEW code.

At the start of training, water-restricted mice were allowed to run on the wheel (Comfort Exercise Wheel, Kaytee Products Inc., Chilton, WI) in their home cage for 3-4 days. The next day they were introduced to the behavioral chamber and were allowed to run on the wheel without reward while photometry data was recorded.

The next day, mice were randomly split into 2 groups – contingent and non-contingent. In the contingent group, reward delivery was contingent on correct behavioral performance, while rewards were independent from behavior for the non-contingent group. Contingent group mice were required to run a minimum of 5 wheel-turns (1000 encoder units) without stopping, and then stop (wheel speed zero) on their own without any external cues. A continuous run of at least this distance followed by a self-initiated stop resulted in 1 second of blue light via the LED and delivery of 7 µl water. Mice collected the reward and then initiated the next trial. If mice stopped before 5 turns, the encoder count was reset, and no reward was delivered. Velocity was calculated every 50 ms from the encoder count. After 7 sessions, an upper bound run distance was introduced. Mice were required to run at least 5 wheel-turns (1000 encoder units) and stop before 9 wheel-turns (1800 encoder units) to get a reward. If they stopped after 9 wheel-turns, the encoder count was reset, and no reward was delivered. The animals then initiated the next trial.

Mice in the non-contingent group did not have any behavioral requirements. They were not required to run and stop to collect the reward, but they typically ran. Every day, the timing of the rewards/cues from a mouse in the contingent group was used to deliver yoked rewards/cues to the non-contingent group of animals. The reward schedules and exposure to the wheel were kept the same for both groups.

#### Passive Approaching Cue/Reward

Animals were placed in a clear plexiglass chamber with a small opening for reward delivery on the left side. The chamber had 2 IR sensors (E18-D80NK, SMAKN, Shenzhen, GD, China), one on each side, and a blue LED on the right side. A linear actuator (L12-100-50-12-P, Actuonix, BC, Canada) was positioned 10 cm away from the clear plexiglass left wall. A custom 3D-printed holder was mounted on a micro linear slide rail (S9-100, Actuonix, BC, Canada), which was connected to the linear actuator. A blue LED and a metal lick spout connected to a custom lickometer were placed on the holder, and the actuator could be programmed to move the LED and the lick spout together to the opening in the left wall. The whole box was placed in a sound-attenuating cubicle (ENV-018MD, Med Associates, Fairfax, VT). The behavioral chamber was controlled using a data acquisition card (USB-6212, National Instruments, Austin, TX) with custom code written in LabVIEW, and behavioral data was acquired by the photometry data acquisition system to allow for synchronized behavioral and neural recording.

Mice were habituated to the behavioral chamber for 1-2 days with a fixed spout and delivery of free reward at constant intervals. After habituation, mice were exposed to the full task. Trials were initiated by breaking the IR beam at the chamber wall closest to the actuator, on the left side. Upon trial initiation the actuator LED illuminated, and the actuator began moving the LED and the reward spout toward the wall opening. The mouse was not required to continuously block the IR sensor during actuator movement. The actuator speed was controlled using a voltage regulator, and it took 8 seconds for the actuator to fully approach. The mouse had full visual access to actuator position at all times because the actuator LED was visible through the clear plexiglass walls. The actuator stopped and the LED extinguished when the reward spout reached the wall opening. Delivery of a 7 µl water reward was triggered by the first lick. If there was no lick within 15 seconds of spout availability, the actuator retracted, and the trial ended. After each trial there was an 8 second inter-trial interval for actuator retraction when a new trial could not be initiated. The end of the inter-trial interval was indicated by LED illumination on the right chamber wall. Mice were required to turn off this LED by blocking the right-side IR sensor before initiating a new trial.

#### Passive Stationary Cue/Reward

Mice were placed in a plastic chamber with a blue LED and a reward spout connected to a custom lickometer. The chamber was located inside a sound-attenuating cubicle (Med Associates, Fairfax, VT). Each trial began with the illumination of a blue LED. After 8 seconds, the LED turned off and a 7 µl water reward was delivered using a variable-rate infusion syringe pump (PHM-107, Med Associates, Fairfax, VT). Licks were detected using a custom lickometer. Inter-trial intervals were drawn from an exponential distribution with a scale parameter of 13 seconds, and a subset of inter-trial intervals between 9 and 40 seconds was used. The behavioral program was controlled using a data acquisition card (USB-6212, National Instruments, Austin, TX) with custom code written in LabVIEW.

### Data analysis

All data analysis and statistical tests were performed using custom code in MATLAB (MathWorks, Natick, MA). The 473 nm and 405 nm fluorescent channels were low-pass filtered at 15 Hz. For the linear track task, the photometry data was then downsampled to 30 Hz using the video TTL pulse times. For all other experiments, the photometry data was not downsampled. The whole-session 405 nm reference channel was fit to the 473 nm channel using linear least squares. Relative fluorescence changes, reported as ΔF/F, were calculated using the following equation: ΔF 473 nm signal-fitted 405 nm signal

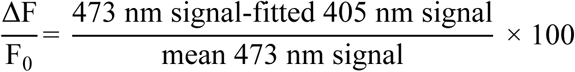

For example traces, the ΔF/F was detrended. For Fig. 1g, h and Extended Data Fig. 3a, b ΔF/F was not detrended or lowpass filtered. For all other analyses, the ΔF/F was lowpass filtered using a Butterworth filter with a cutoff frequency of 0.03 Hz, which was then subtracted from the ΔF/F to remove very slow fluorescence fluctuations.

#### Linear track

The mouse’s position on the track was binned and the mean ΔF/F for each bin was calculated in order to produce position-aligned photometry traces. Trials were averaged for each animal, and population plots were constructed from averages across animals. Speed was processed similarly. For slope calculation, the coefficients were calculated by linearly fitting each trial with polyfit (MATLAB). For fraction of trials with positive slope, the number of trials with positive slope was divided by the total number of trials. Normalized path length was calculated by summing the total distance moved in the x and y directions and normalizing by the length of the track.

#### Running wheel

A run was defined as continuous wheel movement for at least 0.5 wheel-turns (100 encoder counts), and <u>run onset</u> was defined as either the moment this threshold was crossed (with respect to time) or 100 encoder counts (with respect to position). Wheel speed was defined as the distance moved per 50 ms bin measured by counting the number of encoder samples. Each wheel-turn corresponded to a distance of 42 cm. The total encoder count was reset when this number reached zero, which was defined as <u>run offset</u>. Only trials with at least 2.5 wheel-turns (500 encoder counts) were considered for analysis. Trials were rewarded if the total distance was between 5 (1000 encoder count) and 9 (1800 encoder count) wheel-turns. For ΔF/F by position plots, normalized position on each trial ranged from 0 to 1, starting from run onset and stopping at run offset as defined above. ΔF/F traces were averaged for each animal and then averaged across animals for population plots. Slope was calculated for each trial using polyfit (MATLAB). The fraction of trials with positive slope was calculated using only correct trials and was defined as the number of trials with positive slope divided by the total number of trials. For all wheel analyses across sessions, 1 mouse was excluded because of file corruption. Extended Data Fig. 4d, the number of correct trials during the first 50 trials of a session was used to calculate percent correct trials.

#### Approaching and Stationary Conditioned Cue/Reward

Slope was calculated for each trial using data between 1 and 7 seconds following cue/actuator onset. Trials were averaged for each animal and session averages were obtained by averaging across animals. For fraction trials with positive slope, the number of trials with positive slope was divided by the total number of trials.

#### Significance testing, fraction of trials with positive slope

The hierarchical bootstrap method^84^ was used (Fig. 4, Extended Data Table 1). For each session, the number of animals was sampled with replacement, and for each sampled animal trials were sampled with replacement. Then, the proportion of trials with positive slope was calculated. This procedure was repeated 10,000 times. Finally, the probability of the observed fraction of positive slopes given the null hypothesis of an equal fraction of positive and negative slopes was determined from the bootstrap population. This probability was tested for significance with an error rate of 0.05, and Bonferroni correction was used for multiple comparisons.

### Data Availability

The data that support the findings of this study are available from the corresponding author upon reasonable request.

### Code Availability

Code is available from the corresponding author upon reasonable request.

## Acknowledgments

We thank I.T. Ellwood, R.R. Hoy, J.H. Goldberg, J.R. Fetcho, H.K. Reeve, A.K. Churchland, E.L. Troconis, B.J. Sleezer, Y.Y. Ho, W. Gu, Y. Baumel, D.A. Bulkin, R. Chen, A. Roeser, and T. Bollu for helpful discussions; I.T. Ellwood, R.R. Hoy, J.H. Goldberg, A.H. Bass, R.M. Harris-Warrick, C. Linster, N. Yapici, M.J. Sheehan, D.M. Smith, B.J. Sleezer, Y.Y. Ho, and R. Chen for comments on the manuscript; T. Bollu and B.M. Kardon for technical advice and lickometer design; and A.K. Recknagel for technical assistance. This work was supported by the Mong Family Foundation (A.G., C.S.), and by NIH DP2MH109982, the Alfred P. Sloan Foundation, the Whitehall Foundation, and the Brain and Behavior Research Foundation (M.R.W.). M.R.W. is a Robertson Neuroscience Investigator–New York Stem Cell Foundation.

## Author Contributions

A.G. and M.R.W. conceived the project and designed the experiments; A.G. performed stereotaxic surgery; A.G., C.S., R.J.P., D.S.K., and J.A.S. developed behavioral data acquisition methods, conducted photometry experiments and performed histology; C.S. and A.G. developed photometry acquisition methods; A.G. and M.R.W. analysed the data and prepared the manuscript; M.R.W. supervised all aspects of the work.

## Competing Interests

The authors declare no competing interests.

## Additional Information

**Extended Data** is available for this paper.

**Correspondence and requests for materials** should be addressed to M.R.W.

**Extended Data Fig. 1.**
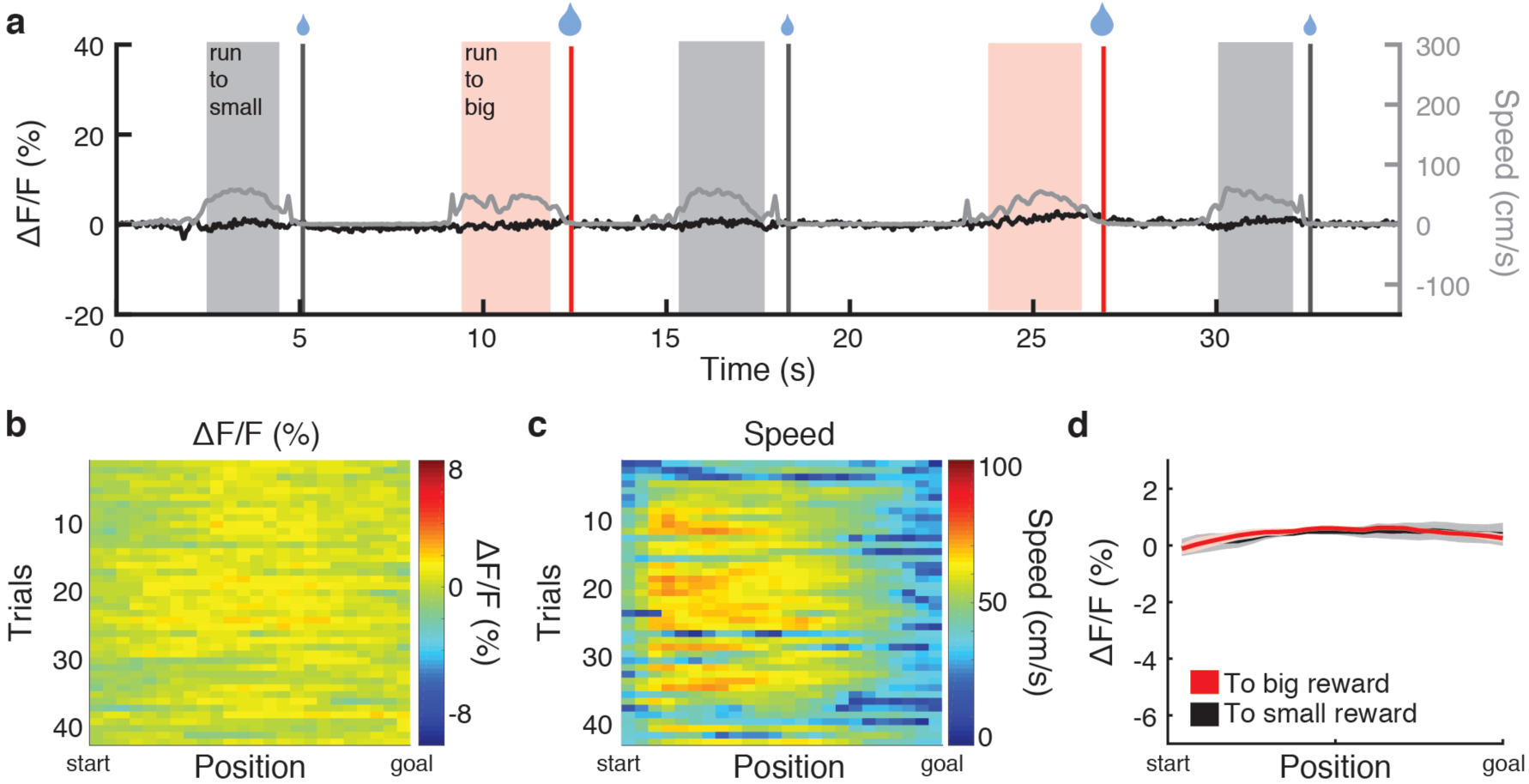
GFP control data. **a**, Example track photometry data from a DAT::GFP mouse. GFP ΔF/F in black, speed in gray. **b**, Example ΔF/F and **c**, speed heatmaps from a DAT::GFP mouse. **d**, Mean GFP ΔF/F with respect to position for big and small rewards (n=3). Error bars indicate s.e.m.

**Extended Data Fig. 2.**
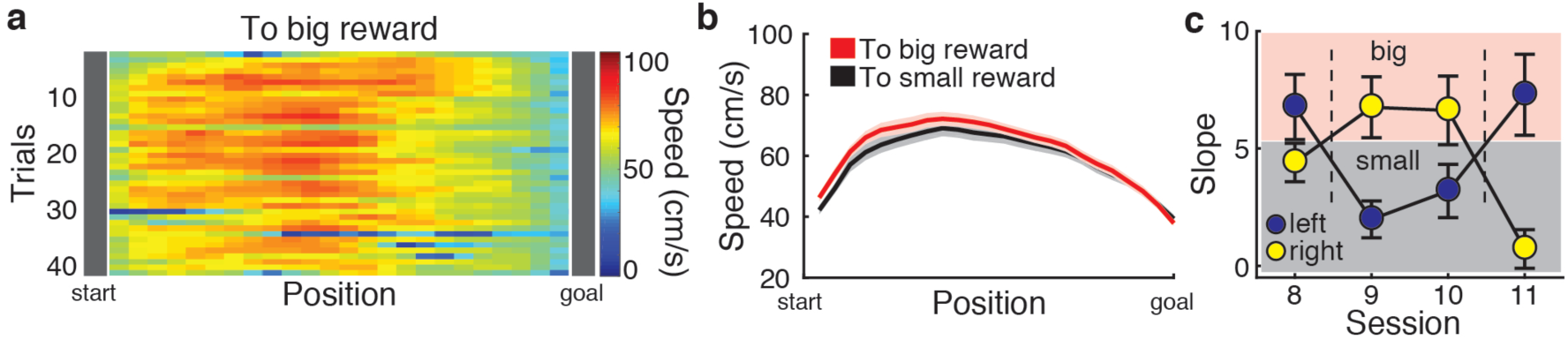
Running speed and reward-switching VTA DA ramp slope data. **a**, Example track speed heatmap from a DAT::GCaMP mouse. **b**, Mean speed during runs to big and small rewards (n=23; 6 animals, 3-4 sessions). **c**, Mean ramp slope following switches in reward position (n=6 for sessions 8, 9, and 11, and n=5 for session 10). Dashed lines indicate switches. Error bars indicate s.e.m.

**Extended Data Fig. 3.**
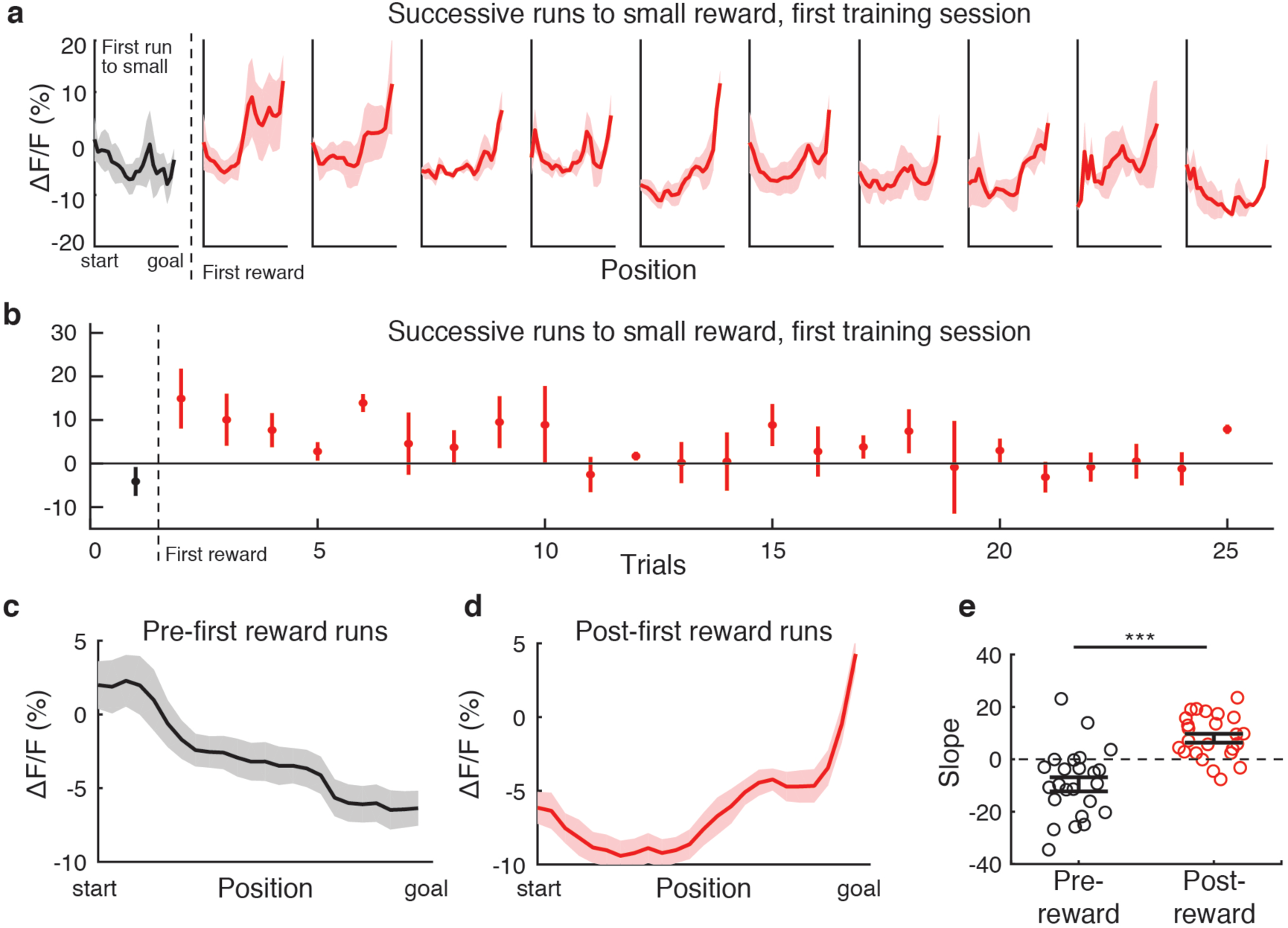
VTA DA ramps during runs to the small reward appear after the first rewarded trial at that location. **a**, Mean ΔF/F for successive runs to the small reward during the first training session. Black – before the first reward at the small goal location; red – after the first reward (n=3). **b**, Ramp slope for successive runs to the small reward during the first track session. **c**, Mean ΔF/F with respect to position for the 8 track runs to the small reward location immediately preceding the first reward delivery (n=24; 3 animals, 8 trials). **d**, Mean ΔF/F with respect to position for the 8 track runs to the small reward location immediately following the first reward delivery (n=24; 3 animals, 8 trials). **e**, Mean ramp slopes for the 8 trials before and after the first reward (n=24; 3 animals, 8 trials). \*\*\**P* < 0.001, Wilcoxon rank-sum test. Error bars indicate s.e.m.

**Extended Data Fig. 4.**
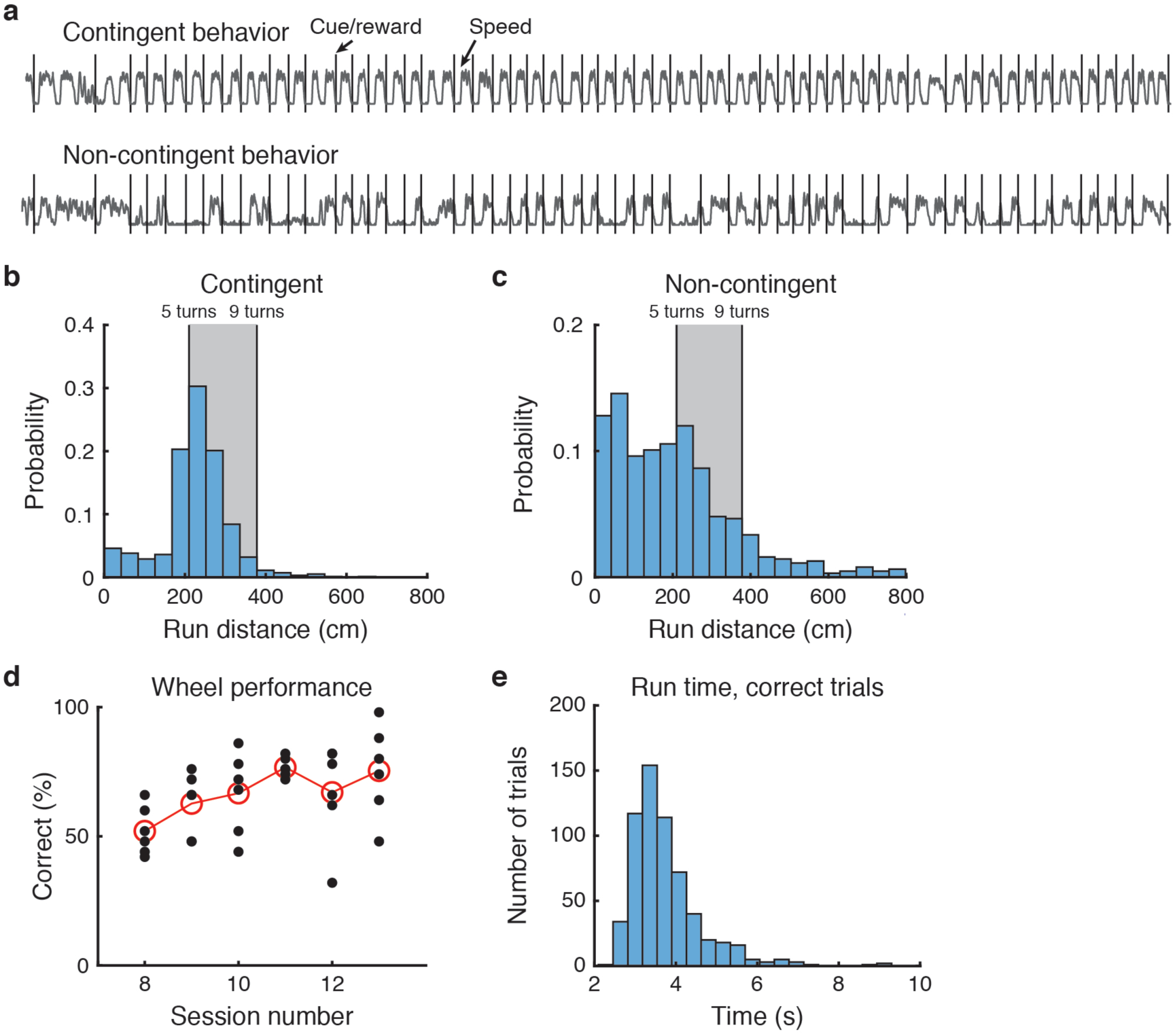
Behavioral performance during wheel running. **a**, Example wheel data from a contingent mouse (top) and a non-contingent mouse (bottom). Cue/reward delivery in black, speed in gray. **b**, Run distance histogram, contingent group. Rewarded distance in gray. **c**, Run distance histogram, non-contingent group. **d**, Wheel performance (percent correct) by session, first 50 trials per session. Mean in red. **e**, Run times for correct runs during the contingent wheel task.

**Extended Data Fig. 5.**
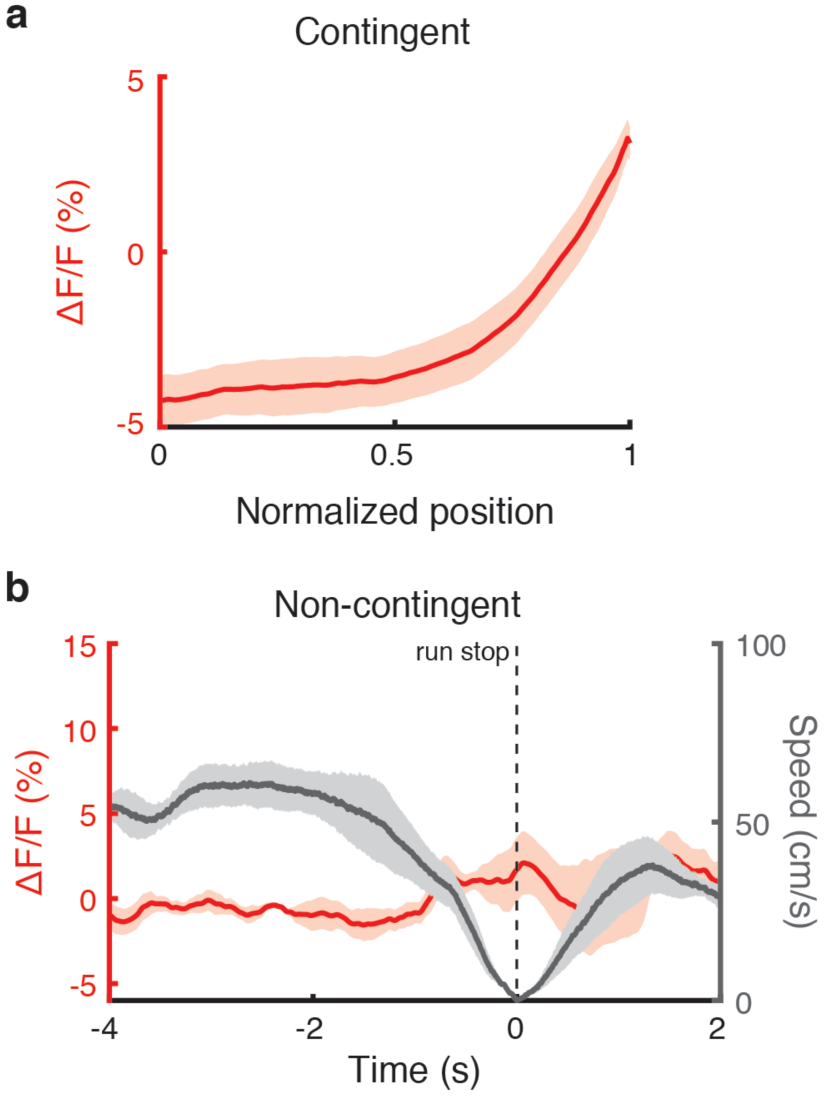
VTA DA neural activity during wheel running. **a**, Mean ΔF/F with respect to position, correct runs, contingent group. **b**, Mean ΔF/F (red) and speed (gray) aligned to stop, runs between 5 and 9 wheel turns with no rewards, non-contingent group. Error bars indicate s.e.m.

**Extended Data Fig. 6.**
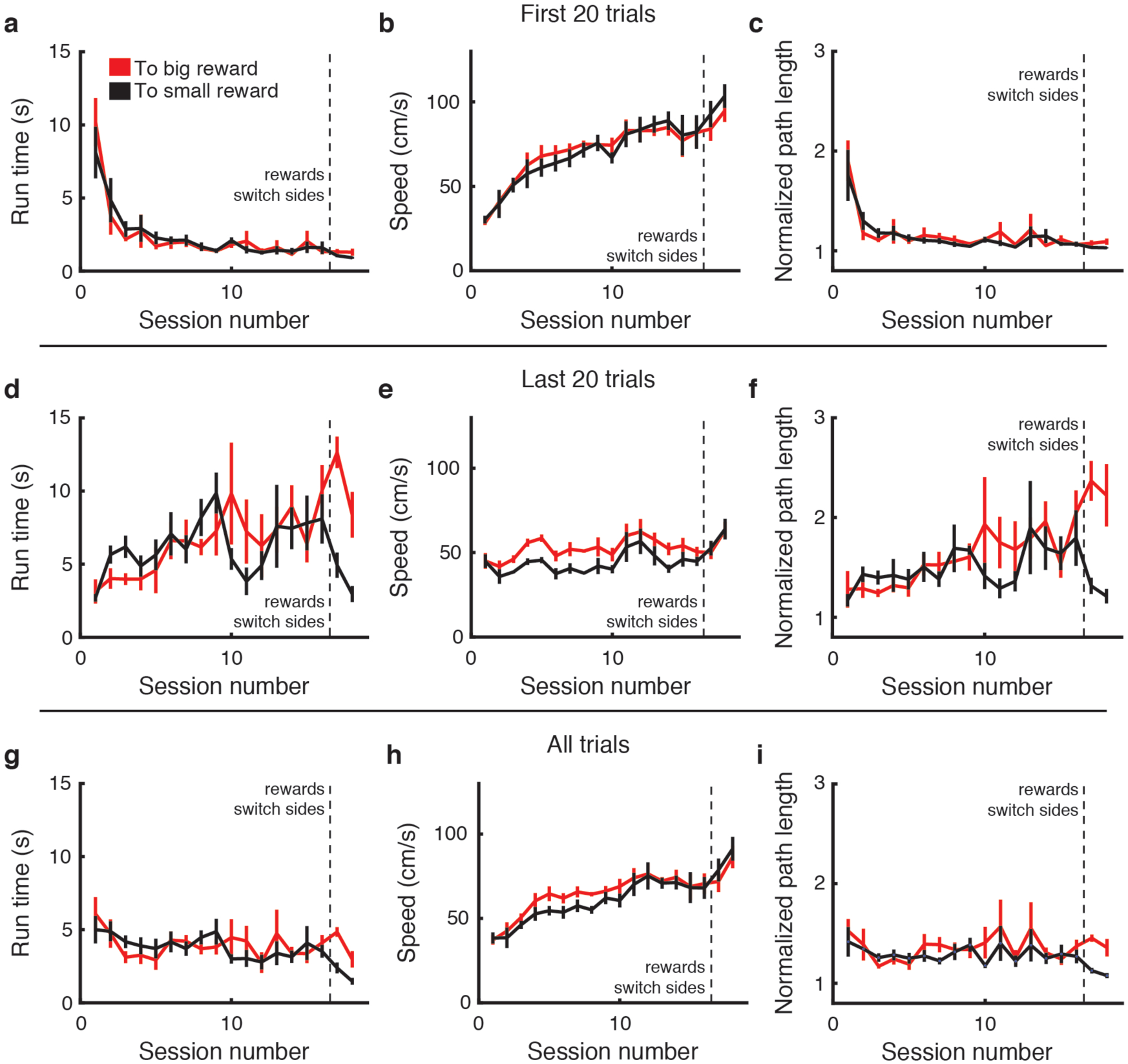
Behavioral performance during navigation toward rewards, track. **a**, Mean run time for track runs to big (red) and small (black) rewards by session, first 20 trials per session (n=3 session 1, n=4 session 2, n=5 sessions 3-18). **b**, Mean speed by session, first 20 trials. **c**, Mean normalized path length by session, first 20 trials. **d, e, f**, Same as a, b, c but using last 20 trials per session. **g, h, i**, Same as a, b, c but using all trials per session. Error bars indicate s.e.m.

**Extended Data Table 1.**
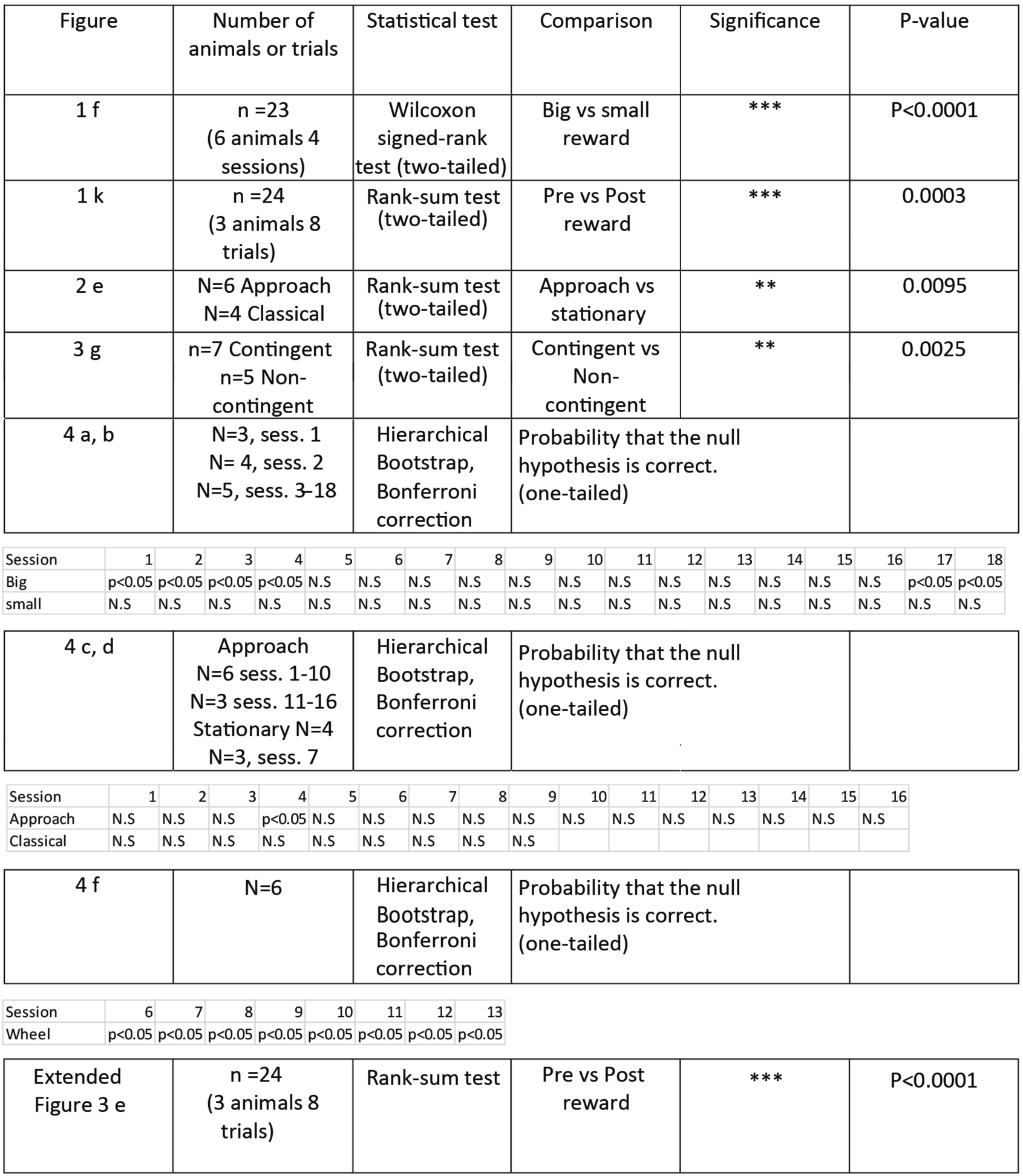
Statistical Results.

| Figure | Number of animals or trials | Statistical test | Comparison | Significance | P-value |
| --- | --- | --- | --- | --- | --- |
| 1 f | n =23<br>(6 animals 4 sessions) | Wilcoxon signed-rank test (two-tailed) | Big vs small reward | *** | P<0.0001 |
| 1 k | n =24<br>(3 animals 8 trials) | Rank-sum test (two-tailed) | Pre vs Post reward | *** | 0.0003 |
| 2 e | N=6 Approach<br>N=4 Classical | Rank-sum test (two-tailed) | Approach vs stationary | ** | 0.0095 |
| 3 g | n=7 Contingent<br>n=5 Non-contingent | Rank-sum test (two-tailed) | Contingent vs Non-contingent | ** | 0.0025 |
| 4 a, b | N=3, sess. 1<br>N= 4, sess. 2<br>N=5, sess. 3-18 | Hierarchical Bootstrap, Bonferroni correction | Probability that the null hypothesis is correct. (one-tailed) |  |  |

| Session | 1 | 2 | 3 | 4 | 5 | 6 | 7 | 8 | 9 | 10 | 11 | 12 | 13 | 14 | 15 | 16 | 17 | 18 |
| --- | --- | --- | --- | --- | --- | --- | --- | --- | --- | --- | --- | --- | --- | --- | --- | --- | --- | --- |
| Big | p<0.05 | p<0.05 | p<0.05 | p<0.05 | N.S | N.S | N.S | N.S | N.S | N.S | N.S | N.S | N.S | N.S | N.S | N.S | p<0.05 | p<0.05 |
| small | N.S | N.S | N.S | N.S | N.S | N.S | N.S | N.S | N.S | N.S | N.S | N.S | N.S | N.S | N.S | N.S | N.S | N.S |

|  |  |  |  |
| --- | --- | --- | --- |
| 4 c, d | Approach<br>N=6 sess. 1-10<br>N=3 sess. 11-16<br>Stationary N=4<br>N=3, sess. 7 | Hierarchical Bootstrap, Bonferroni correction | Probability that the null hypothesis is correct. (one-tailed) |

| Session | 1 | 2 | 3 | 4 | 5 | 6 | 7 | 8 | 9 | 10 | 11 | 12 | 13 | 14 | 15 | 16 |
| --- | --- | --- | --- | --- | --- | --- | --- | --- | --- | --- | --- | --- | --- | --- | --- | --- |
| Approach | N.S | N.S | N.S | p<0.05 | N.S | N.S | N.S | N.S | N.S | N.S | N.S | N.S | N.S | N.S | N.S | N.S |
| Classical | N.S | N.S | N.S | N.S | N.S | N.S | N.S | N.S | N.S |  |  |  |  |  |  |  |

|  |  |  |  |
| --- | --- | --- | --- |
| 4 f | N=6 | Hierarchical Bootstrap, Bonferroni correction | Probability that the null hypothesis is correct. (one-tailed) |

| Session | 6 | 7 | 8 | 9 | 10 | 11 | 12 | 13 |
| --- | --- | --- | --- | --- | --- | --- | --- | --- |
| Wheel | p<0.05 | p<0.05 | p<0.05 | p<0.05 | p<0.05 | p<0.05 | p<0.05 | p<0.05 |

2 Extended Data Table 1 | Statistical Results.
|  |  |  |  |  |  |
| --- | --- | --- | --- | --- | --- |
| Extended Figure 3 e | n =24<br>(3 animals 8 trials) | Rank-sum test | Pre vs Post reward | *** | P<0.0001 |

## Notes

### Competing Interest Statement

The authors have declared no competing interest.

